# 5-ASA can functionally replace Clostridia to prevent a post-antibiotic bloom of *Candida albicans* by maintaining epithelial hypoxia

**DOI:** 10.1101/2023.04.17.537218

**Authors:** Hannah P. Savage, Derek J. Bays, Mariela A. F. Gonzalez, Eli. J. Bejarano, Henry Nguyen, Hugo L. P. Masson, Thaynara P. Carvalho, Renato L. Santos, George R. Thompson, Andreas J. Bäumler

## Abstract

Antibiotic prophylaxis sets the stage for an intestinal bloom of *Candida albicans*, which can progress to invasive candidiasis in patients with hematologic malignancies. Commensal bacteria can reestablish microbiota-mediated colonization resistance after completion of antibiotic therapy, but they cannot engraft during antibiotic prophylaxis. Here we use a mouse model to provide a proof of concept for an alternative approach, which replaces commensal bacteria functionally with drugs to restore colonization resistance against *C. albicans*. Streptomycin treatment, which depletes Clostridia from the gut microbiota, disrupted colonization resistance against *C. albicans* and increased epithelial oxygenation in the large intestine. Inoculating mice with a defined community of commensal Clostridia species reestablished colonization resistance and restored epithelial hypoxia. Notably, these functions of commensal Clostridia species could be replaced functionally with the drug 5-aminosalicylic acid (5-ASA), which activates mitochondrial oxygen consumption in the epithelium of the large intestine. When streptomycin-treated mice received 5-ASA, the drug reestablished colonization resistance against *C. albicans* and restored physiological hypoxia in the epithelium of the large intestine. We conclude that 5-ASA treatment is a non-biotic intervention that restores colonization resistance against *C. albicans* without requiring the administration of live bacteria.

## INTRODUCTION

Invasive candidiasis, defined as *Candida* spp. in the blood (candidemia) and deep-seated infections including hepatosplenic candidiasis, is a leading cause of nosocomial infections in the United States, particularly among those with hematologic malignancies (Wilson, Reyes et al. 2002, Magill, O’Leary et al. 2018). Invasive candidiasis carries a high mortality rate (up to 49%) and places a large financial burden on our healthcare system (Viscoli, Girmenia et al. 1999, Wilson, Reyes et al. 2002, Gudlaugsson, Gillespie et al. 2003, Morgan, Meltzer et al. 2005). Therefore, there is a critical need for novel approaches for preventing or treating invasive candidiasis.

*Candida albicans* is the most common causative species of invasive candidiasis and has been found to colonize the human gastrointestinal (GI) tract as a commensal component of the microbiome in approximately 60% of humans (Pappas, Lionakis et al. 2018). *Candida* spp. Have the ability to form biofilms on prosthetic devices such as indwelling venous catheters leading to candidemia (Pappas, Lionakis et al. 2018), however, it is thought that dissemination from the GI tract is the most common etiology of invasive disease in patients with hematologic malignancies. Invasion of *C. albicans* from the GI tract is a consequence of hematologic malignancy treatment, which induces neutropenia and GI epithelial disruption (Koh, Kohler et al. 2008). Notably, *Candida* spp. expand in the GI tract prior to translocating into the blood stream in patients with hematologic malignancy undergoing stem cell transplants (Zhai, Ola et al. 2020, Alonso-Monge, Gresnigt et al. 2021). This bloom of *Candida* spp. in the GI tract is triggered by antibiotics, which disrupt the gut microbiota to promote growth of the opportunistic pathogen (Pappas, Lionakis et al. 2018, Zhai, Ola et al. 2020). As invasive candidiasis originates from the gut, preventing an expansion of intestinal *Candida* spp. during antibiotic prophylaxis could be a novel approach to reducing invasive disease in patients with hematologic malignancy. But developing this prophylactic strategy requires a better understanding of the factors promoting and protecting against a bloom of *Candida* spp. in the GI tract.

The gut microbiota is dominated by obligately anaerobic bacteria belonging to the classes Clostridia and Bacteroidia (Human Microbiome Project 2012). The importance of an intact microbiota in preventing an expansion of *C. albicans* in the gastrointestinal tract can be modeled in mice (Clark 1971, Kennedy and Volz 1985, Samonis, Anastassiadou et al. 1994, Mason, Erb Downward et al. 2012). A disruption of the gut microbiota, particularly through administration of broad-spectrum antibiotics with anaerobic activity, drives an intestinal *C. albicans* bloom, suggesting that obligately anaerobic bacteria confer colonization resistance against this opportunistic pathogen in mice (Kennedy and Volz 1985, Samonis, Anastassiadou et al. 1994, Mason, Erb Downward et al. 2012, Gutierrez, Weinstock et al. 2020). It has been proposed that colonization resistance is due to the production of metabolites by obligately anaerobic bacteria that inhibit growth of *C. albicans*. For instance, members of the class Clostridia are the main producers of the short-chain fatty acid butyrate in the gut microbiota (Louis and Flint 2009, Vital, Howe et al. 2014), a metabolite that inhibits growth of *C. albicans in vitro* (Nguyen, Lopes et al. 2011). An alternative hypothesis is that Clostridia confer colonization resistance by influencing epithelial physiology. For example, Clostridia induce the epithelial release of antimicrobial peptides, which correlates with reduced gut colonization by *C. albicans* (Fan, Coughlin et al. 2015). Furthermore, Clostridia-derived butyrate activates the nuclear receptor PPAR-γ (peroxisome proliferator-activated receptor gamma) in the colonic epithelium to increase mitochondrial oxygen consumption, which maintains the mucosal surface in a state of physiological hypoxia (<1% O_2_) to maintain anaerobiosis (Byndloss, Olsan et al. 2017). Antibiotic treatment depletes butyrate (Meynell 1963), resulting in a metabolic reprogramming of the epithelium to increase epithelial oxygenation (Byndloss, Olsan et al. 2017). As a result, oxygen diffuses into lumen of the of the antibiotic-treated colon to disrupt anaerobiosis, thereby weakening colonization resistance against facultatively anaerobic bacteria, such as *Enterobacteriaceae* (Rivera-Chavez, Zhang et al. 2016, Byndloss, Olsan et al. 2017). It remains to be determined which of these mechanisms is causatively linked to a loss of colonization resistance against *C. albicans* during antibiotic prophylaxis *in vivo*.

Antibiotic prophylaxis is a necessary treatment for many patients with hematologic malignancies (Moreno-Sanchez and Gomez-Gomez 2022), which limits the options available for restoring normal microbiota function in this patient population. For example, it is not possible to use inoculation with Clostridia species to restore colonization resistance against *C. albicans* until antibiotic prophylaxis has been discontinued, but patients often develop invasive candidiasis while they are still on antibiotics. The hypothesis that Clostridia confer colonization resistance by influencing epithelial physiology suggests that it should be possible to develop drugs that target the epithelium to restore colonization resistance against *C. albicans* by replacing the microbiota functionally. This approach is appealing because drugs can be applied concurrently with antibiotic prophylaxis. The goal of this study was to explore the feasibility of this approach.

## RESULTS

### Clostridia confer colonization resistance against *C. albicans*

We first sought to establish a mouse model of increased susceptibility to *Candida albicans* colonization due to microbiota changes after antibiotic treatment. Pre-treatment with a single dose of streptomycin decreased colonization resistance to a subsequent *C. albicans* challenge in specific pathogen-free C57BL/6J mice by more than 2 logs of exposure (Fig. 1A), which is consistent with prior findings (Gutierrez, Weinstock et al. 2020). Similarly, in mice pre-colonized with *C. albicans*, treatment with a single dose of streptomycin triggered a fecal bloom of the opportunistic pathogen (Fig. 1B). To confirm that the increased colonization is due to changes in the microbiota and not direct effects of the antibiotic on *C. albicans* or the host cells, we colonized germ-free Swiss Webster mice with cecal contents (cecal microbiota transfer, CMT) from C57BL/6J mice collected 48 hours after mock-treatment or streptomycin treatment (Fig. 2A). The CMT from antibiotic-treated mice provided reduced colonization resistance compared to the CMT from control mice when gnotobiotic mice were challenged with *C. albicans* (Fig. 2B). To confirm that reduced colonization resistance was due to loss of bacteria after antibiotic treatment, germ-free mice engrafted an CMT from antibiotic-treated mice received a second CMT from mock-treated mice one week later, which restored colonization resistance against *C. albicans* challenge (Fig. 2B).

**Figure 1:**
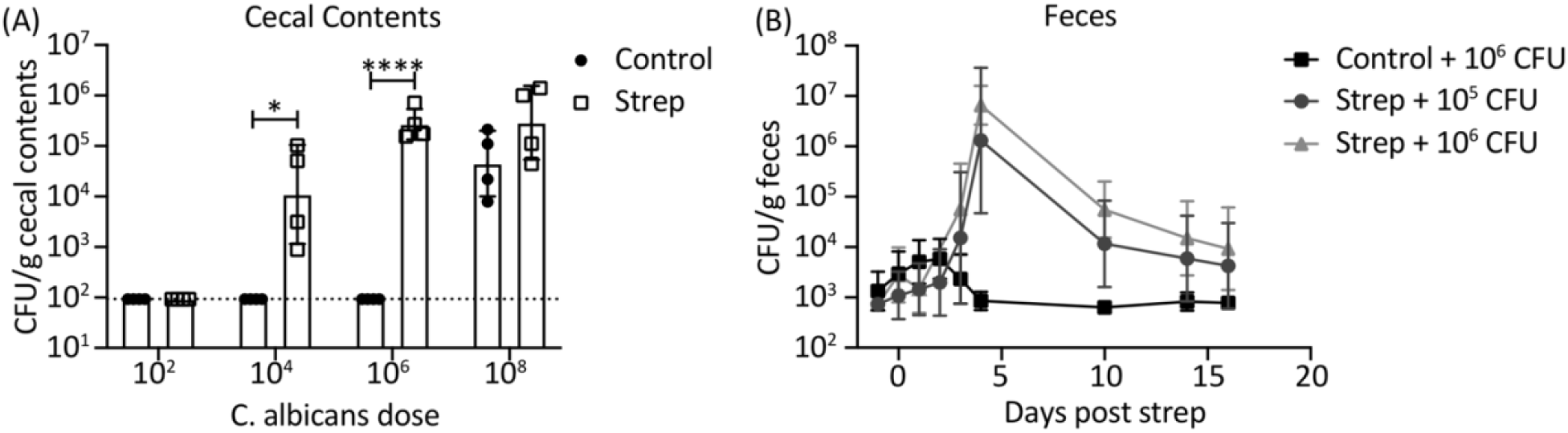
Antibiotic treatment increases susceptibility to *Candida albicans* colonization in mice. (A) SPF C57BL/6J mice were given 20 mg streptomycin (strep) as a single oral gavage or mock treated (Control). Mice were infected with the indicated dose of *C. albicans* 48 hours later. Shown are colony-forming units (CFU) of *C. albicans*/g (symbols) and geometric mean +/-geometric SD (bar) recovered from cecal contents of Control or Strep-treated mice 7 days after infection (n=4). (B) SPF C57BL/6 mice were infected with the indicated dose of *C. albicans* and given 20 mg streptomycin (strep) by oral gavage five days after infection. Shown are geometric mean +/-geometric SD of the CFU *C. albicans*/g recovered from feces (1-2 pellets per mouse) of Control or Strep-treated mice at the indicated time points (n=4 for Control, n=8 for Strep-treated). *, *P* < 0.05; ****, *P* < 0.00005.

**Figure 2:**
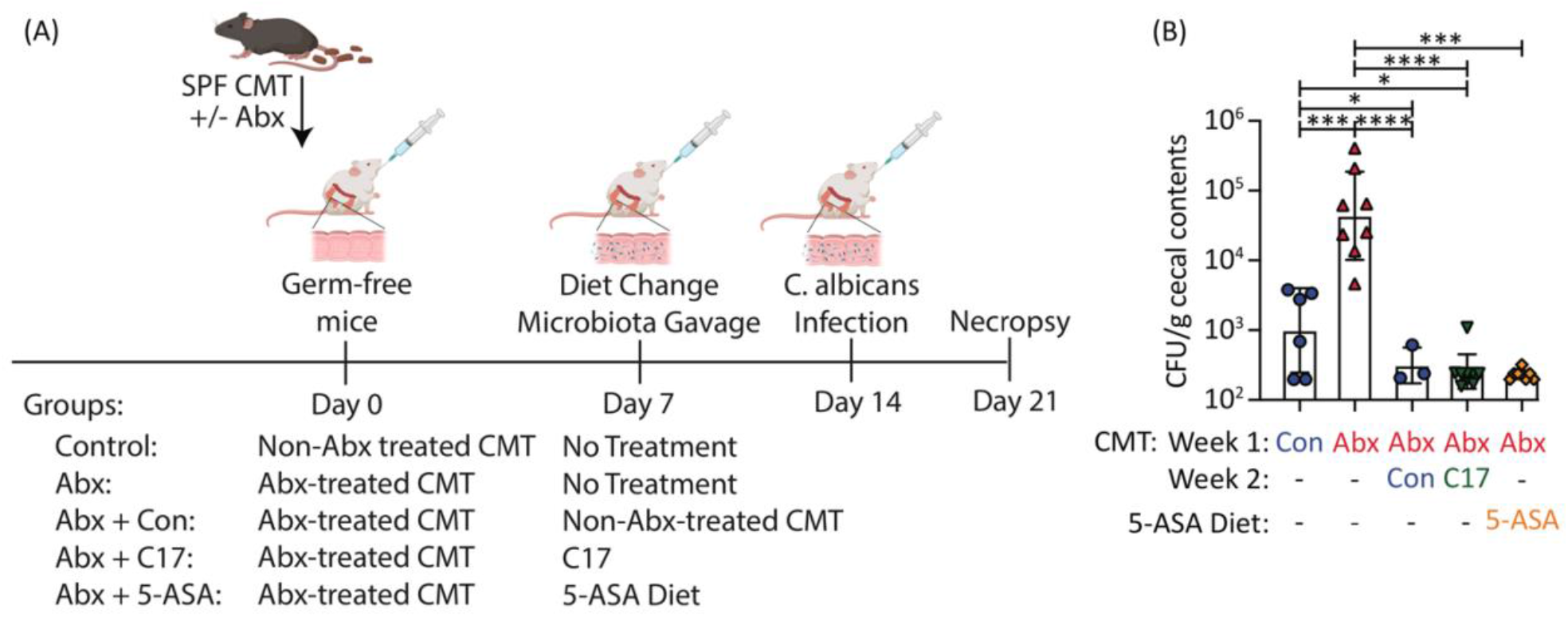
Clostridia and 5-ASA provide colonization resistance against *C. albicans*. (A) Germ-free Swiss Webster mice were engrafted with a cecal microbiota transplant (CMT) from C57BL/6J (Con in panel B) mice (Non-Abx-treated CMT in panel A) and streptomycin-treated (Abx in panel B) C57BL/6J mice (Abx treated CMT in panel A). As indicated, groups of mice received a second Non-Abx-treated CMT, a community of 17 human Clostridia isolates (C17) or were started on 5-ASA-containing chow after 7 days. Mice were infected with *C. albicans* after an additional 7 days, as indicated. (B) Shown are colony-forming units (CFU) of *C. albicans* (each symbol represents data from one animal) and geometric mean +/-geometric SD (bar) recovered from cecal contents of each group 7 days after infection (n=3 for Abx + Con, n= 6 for Controls, n=8-9 for remaining groups). *, *P* < 0.05; ***, *P* < 0.0005; ****, *P* < 0.00005. Figure 2A was created with BioRender.com.

We have previously shown that a single dose of streptomycin depletes Clostridia from the gut microbiota to lower colonization resistance against *Enterobacteriaceae* (Byndloss, Olsan et al. 2017). To investigate whether members of the order Clostridia are responsible for loss of colonization resistance against *C. albicans* after streptomycin treatment, germ-free mice engrafted with a CMT from streptomycin-treated C57BL/6J mice were inoculated with a community of 17 human Clostridia isolates (C17), which induce the formation of regulatory T cells in the colonic mucosa of gnotobiotic mice (Atarashi, Tanoue et al. 2013) and produce the short-chain fatty acids acetate and butyrate (Narushima, Sugiura et al. 2014). Inoculation with this consortium of 17 human Clostridia isolates restored colonization resistance against *C. albicans* in germ-free mice engrafted with a CMT from streptomycin-treated C57BL/6J mice (Fig. 2B). Collectively, these data suggested that streptomycin treatment weakens colonization resistance against *C. albicans* by depleting Clostridia from the gut microbiota.

### The drug 5-aminosalicylic acid restores colonization resistance against *C. albicans* after streptomycin treatment

Next, we wanted to determine whether drugs could replace Clostridia species functionally to restore colonization resistance against *C. albicans* after streptomycin treatment. Conventional wisdom holds that Clostridia species produce short chain fatty acids, which reduce growth (Nguyen, Lopes et al. 2011, Guinan, Wang et al. 2019) and hyphal formation by *C. albicans in vitro (Noverr and Huffnagle 2004)*, but the *in vivo* evidence supporting these inhibitory activities of short-chain fatty acids remains correlative. *In vivo*, Clostridia-derived butyrate activates PPAR-γ signaling in the colonic epithelium to enhance mitochondrial oxygen consumption, which is important for maintaining anaerobiosis (Byndloss, Olsan et al. 2017). We have shown previously that the latter activity of Clostridia-derived butyrate can be replaced with the PPAR-γ agonist 5-aminosalicylic acid (5-ASA), a drug that restores an anaerobic environment in the colon of mice with chemically-induced colitis to limit growth of facultatively anaerobic *Enterobacteriaceae* (Cevallos, Lee et al. 2021). Since *C. albicans* is also a facultatively anaerobic microbe, we reasoned that a disruption of anaerobiosis during streptomycin treatment (Rivera-Chavez, Zhang et al. 2016, Byndloss, Olsan et al. 2017) might enhance growth of the opportunistic pathogen, but this idea has never been tested. This hypothesis would imply that restoration of anaerobiosis using 5-ASA could prevent the antibiotic-induced expansion of the facultatively anaerobic fungus *Candida* by restoring epithelial hypoxia.

To investigate whether a direct inhibition of *C. albicans* by butyrate or butyrate-mediated PPAR-γ activation were required for colonization resistance against *C. albicans*, we treated mice with 5-ASA, which activates PPAR-γ signaling specifically in the intestinal epithelium (Cevallos, Lee et al. 2021). C57BL/6J mice received normal chow or chow supplemented with 5-ASA and were mock-treated or pre-treated with a single dose of streptomycin to decrease colonization resistance. Remarkably, 5-ASA supplementation protected streptomycin pre-treated mice from a subsequent *C. albicans* challenge, as indicated by reduced recovery of the opportunistic pathogen from cecal contents assessed at one (Fig. 3A) or seven (Fig. 3B) days after challenge. However, 5-ASA supplementation did not restore butyrate concentrations in the feces of streptomycin pre-treated mice (Fig. 4). Similarly, in mice pre-colonized with *C. albicans*, treatment with a single dose of streptomycin triggered a fecal bloom of the opportunistic pathogen, which was blunted in mice receiving 5-ASA supplementation (Fig. 3C). Finally, 5-ASA supplementation restored colonization resistance against *C. albicans* in germ-free mice engrafted a CMT from antibiotic-treated mice (Fig. 2B). Collectively, these data suggested that treatment with 5-ASA restores colonization resistance against *C. albicans* after streptomycin treatment.

**Figure 3:**
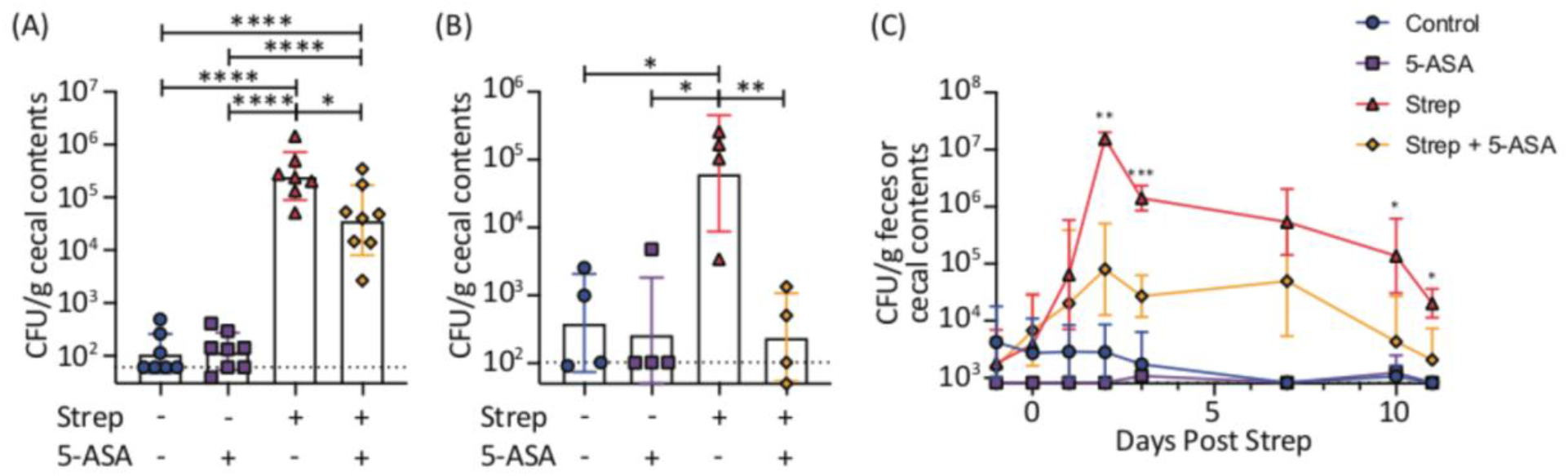
5-ASA restores colonization resistance against *C. albicans* after antibiotic treatment. (A-B) Mice were started on 5-ASA-containing chow 2 days prior to streptomycin (strep) treatment (20 mg strep by oral gavage), and infected 1-2 days after strep with (A) 10^5^ or (B) 10^6^ colony-forming units (CFU) of *C. albicans*. Shown are CFU *C. albicans*/g (each symbol represents data from one animal) and geometric mean +/-geometric SD (bar) recovered from cecal contents of *C. albicans*-colonized mice on (A) day 1 (n=7-8) or (B) day 7 (n=4) after infection. (C) Mice were started on a 5-ASA-containing chow and given 10^6^ CFU *C. albicans*, colonization was confirmed, and mice were given 20 mg streptomycin (strep) by oral gavage 3 days later. Shown are geometric mean +/-geometric SD of the CFU *C. albicans*/g recovered from feces (1-2 pellets per mouse) of each group at the indicated time points (n=6 for strep + 5-ASA, n=3 for remaining groups). *, *P* < 0.05; **, *P* < 0.005; ***, *P* < 0.0005; ****, *P* < 0.00005.

**Figure 4:**
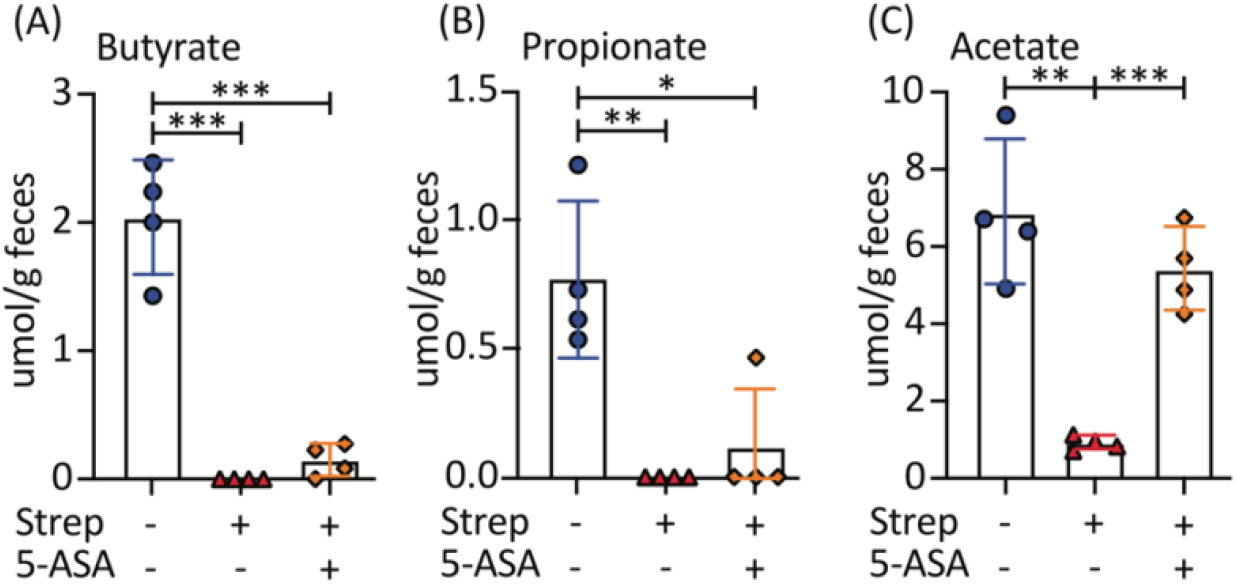
5-ASA does not restore butyrate after an antibiotic-mediated loss of short chain fatty acids. Shown are concentrations (μmol/g feces) +/-SD of (A) butyrate, (B) propionate, and (A) acetate in the feces of individual mice (each symbol represents data from one animal). Antibiotic-treated mice (Strep) were given 20 mg streptomycin by oral gavage one day prior to feces collection. Mice that received 5-ASA (Strep + 5-ASA) were placed on a 5-ASA-containing chow 5 days prior to treatment with streptomycin (n=4). *, *P* < 0.05; **, *P* < 0.005; ***, *P* < 0.0005.

### 5-ASA treatment restores colonization resistance by restoring epithelial hypoxia

5-ASA and butyrate both have anti-inflammatory effects (Segain, Raingeard de la Blétière et al. 2000, Ricote and Glass 2007), and inflammation can alter the microbial composition of the gut, resulting in a bloom of facultatively anaerobic bacteria (Winter, Winter et al. 2013). To investigate whether the anti-inflammatory effects of 5-ASA explain the reduced colonization by *C. albicans*, we investigated whether streptomycin treatment or *C. albicans* challenge trigger inflammatory responses in the gut. Limited to no inflammation was observed in germ-free mice engrafted with CMT from streptomycin-treated C57BL/6J mice during *C. albicans* challenge as indicated by histopathological scoring of sections from the cecum or colon (Fig. 5A-B). In streptomycin pre-treated C57BL/6J mice, transcript levels of genes encoding lipocalin-2, TNF-α, or IL-17A were measured one day after challenge with *C. albicans* (Fig. 5C). Although a small increase in transcript levels of *Lcn2* and *Il17a* was observed in streptomycin treatment during *C. albicans* challenge, these increases were not reversed by 5-ASA treatment and thus did not correlate with the reduction in *C. albicans* colonization seen in these animals (Fig. 3A). In summary, these data did not support the idea that the anti-inflammatory activity of 5-ASA is responsible for its ability to restore colonization resistance against *C. albicans*.

**Figure 5:**
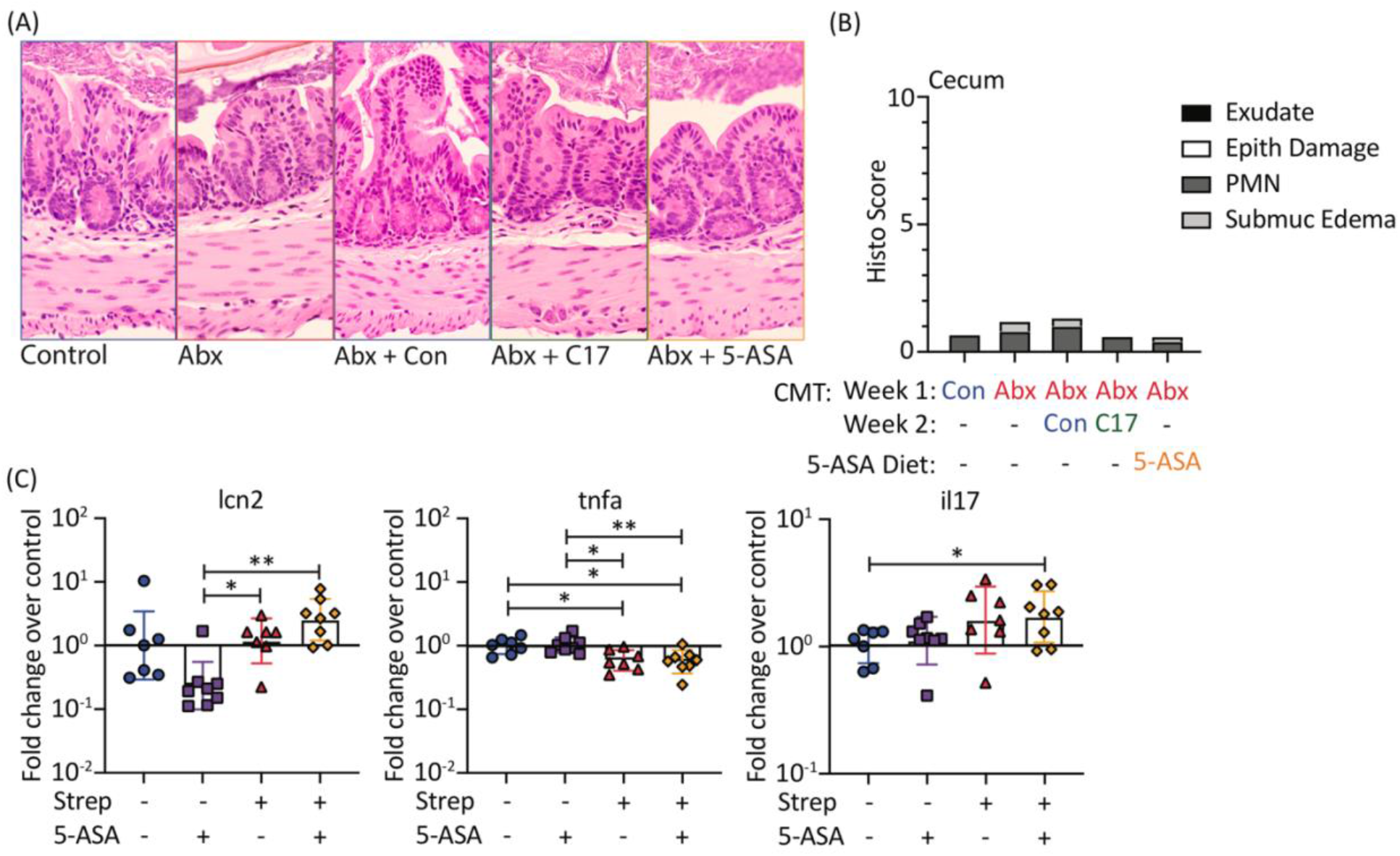
*C. albicans* challenge is not associated with overt intestinal inflammation. Shown are (A) representative images and (B) mean histopathology scores of the ceca of germ-free mice engrafted with a cecal microbiota transplant (CMT) from C57BL/6J (Control) mice or streptomycin-treated (Abx). Seven days later, the latter group received a second CMT from C57BL/6J mice (Abx + Con), a community of 17 human Clostridia isolates (Abx + C17) or were started on 5-ASA-containing chow (Abx + 5-ASA). Mice were infected with *C. albicans* after an additional 7 days. Histological sections were blinded and scored by a veterinary pathologist. (C) Mice were started on 5-ASA-containing chow 2 days prior to streptomycin (strep) treatment (20 mg strep by oral gavage), and infected 1 day after strep with 10^5^ colony-forming units (CFU) of *C. albicans*. Colonocytes were collected one day after infection to extract RNA for quantitative real-time PCR of pro-inflammatory markers. Shown are fold change in expression of the indicated genes (*Lcn2, Tnfa, Il17a*) in colonocytes of individual mice (each symbol represents data from one animal) compared to the relative expression in the control group (geometric mean) and the geometric mean +/-geometric SD (bar) of each group (n= 7-8). *, *P* < 0.05; **, *P* < 0.005.

We have previously shown that during homeostasis in healthy colons, butyrate activates PPAR-γ in host colonocytes resulting in epithelial hypoxia, which limits diffusion of oxygen into the colonic lumen (Byndloss, Olsan et al. 2017). *Candida* spp. respire oxygen, so an increase in epithelial oxygenation triggered by antibiotic-mediated Clostridia depletion might enhance growth of this facultatively anaerobic opportunistic pathogen. To test this idea, we visualized epithelial oxygenation with pimonidazole, a 2-nitroimidazole that is reductively activated specifically in hypoxic cells (< 1% oxygen) (Kizaka-Kondoh and Konse-Nagasawa 2009, Rijk, van Schaik et al. 1988). Pimonidazole staining revealed that epithelial oxygenation was increased in both antibiotic-treated C57BL/6J mice (Fig. 6A-C) and germ-free Swiss Webster mice engrafted with a CMT from antibiotic-treated C57BL/6J mice (Fig. 6D-F), compared to antibiotic naïve controls. Inoculation with a consortium of 17 human Clostridia isolates restored epithelial in germ-free mice engrafted with a CMT from streptomycin-treated C57BL/6J mice (Fig. 6D-F). These data confirmed that Clostridia function in maintaining epithelial hypoxia in the colon (Byndloss, Olsan et al. 2017). Importantly, treatment with 5-ASA restored epithelial hypoxia in antibiotic-treated C57BL/6J mice (Fig. 6A-C) and in germ-free mice engrafted with a CMT from antibiotic-treated C57BL/6J mice (Fig. 6D-F). These results support the idea that role of Clostridia in maintaining epithelial hypoxia in the large intestine can be functionally replaced by the drug 5-ASA.

**Figure 6:**
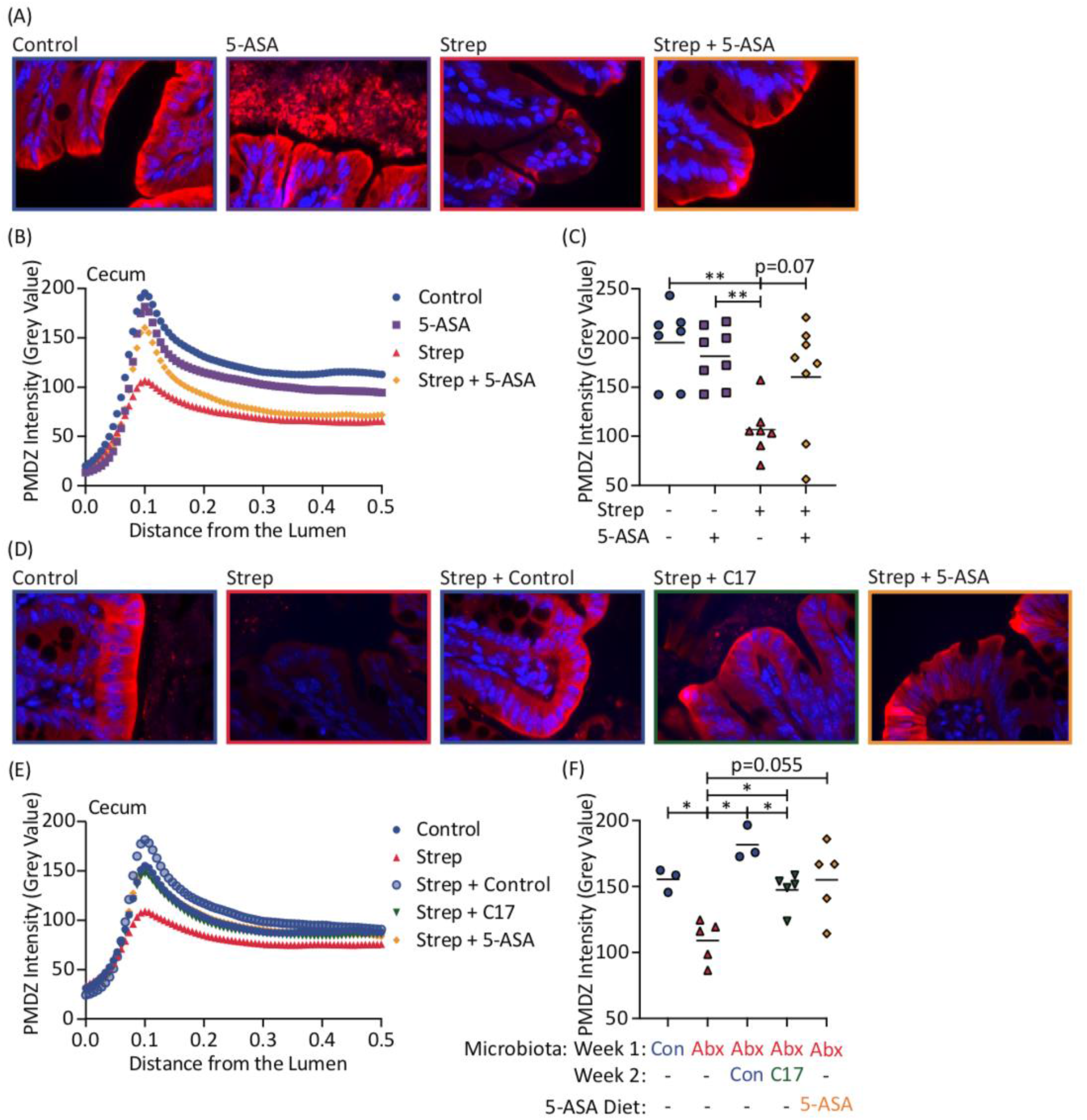
Epithelial hypoxia is decreased by streptomycin treatment but increased by 5-ASA. (A-C) Mice were started on 5-ASA-containing chow 2 days prior to streptomycin (strep) treatment (20 mg strep by oral gavage), and infected 1 day after strep with 10^5^ colony-forming units (CFU) of *C. albicans*. Cecal tips were collected on Day 1 post-infection. 30-90 minutes prior to necropsy, mice were injected with pimonidazole (PMDZ). Samples were stained with mouse anti-PMDZ monoclonal antibody and cyanine3-labeled goat anti-mouse IgG to detect PMDZ binding to hypoxic tissues. Shown are (A) representative images of samples stained for PMDZ (red) and counterstained with DAPI (blue); (B) mean PMDZ intensities of each group from the lumen (distance of 0.0 arbitrary units) to the border of the colonocytes (distance of 0.1), and into the tissue; and (C) peak PMDZ intensity for each mouse (each symbols represents data from one animal) and mean peak intensity for each group (lines) (n=7-8). (D-F) Germ-free mice were engrafted with a cecal microbiota transplant (CMT) from C57BL/6J (Control) mice or streptomycin-treated (Abx). Seven days later, the latter group received a second CMT from C57BL/6J mice (Abx + Con), a community of 17 human Clostridia isolates (Abx + C17) or were started on 5-ASA-containing chow (Abx + 5-ASA). Mice were infected with *C. albicans* after an additional 7 days. Cecal tips were collected on Day 7 post-infection. 30-90 minutes prior to necropsy, mice were injected with pimonidazole (PMDZ). Shown are (D) representative images of samples stained for PMDZ (red) and DAPI (blue); (E) mean PMDZ intensities of each group from the lumen (distance of 0.0 arbitrary units) to the border of the colonocytes (distance of 0.1), and into the tissue; and (F) peak PMDZ intensity for each mouse (symbols) and mean peak intensity for each group (lines) (n=3 for Control and Abx+Con, n=5 for remaining groups). *, *P* < 0.05; **, *P* < 0.005; ***, *P* < 0.0005; ****, *P* < 0.00005.

## DISCUSSION

Our results support the idea that Clostridia contribute to colonization resistance against

*C. albicans*. Thus, direct restoration of butyrate-producing Clostridia could be considered a promising probiotic therapy to restore colonization resistance in patients at risk for developing invasive candidiasis. However, as the population greatest at risk of invasive candidiasis are those with hematologic malignancies, this strategy raises safety concerns. While probiotics can act as PPAR-γ agonists (Hasan, Rahman et al. 2019) and reduce Candida colonization (Archambault and Dongari-Bagtzoglou 2022), probiotics or full fecal microbiota transplants are not typically considered in hematologic malignancy due to safety concerns over infections stemming from the deleterious effects of chemotherapy (Ma, Li et al. 2021). Additionally, in patients on antibiotic prophylaxis, probiotics or fecal microbiota transplants are not likely to engraft successfully.

Here we demonstrate for the first time that post-antibiotic *Candida albicans* expansion can be restricted by supporting colonic epithelial hypoxia through activation of the PPAR-γ pathway. The *in vivo* inhibition of an *C. albicans* expansion by Clostridia*-*derived butyrate or 5-ASA is due to restoration of epithelial hypoxia, rather than through direct interactions with *C. albicans in vivo* or PPAR-γ’s anti-inflammatory effects. To our knowledge, this is the first time that 5-ASA efficacy has been documented in a non-inflammatory environment to prevent the expansion of a facultative anaerobic opportunistic pathogen. Although typically considered an anti-inflammatory drug, we show that 5-ASA was also able to promote healthy colonocyte metabolism to maintain hypoxia in the colon after antibiotic-induced non-inflammatory dysbiosis. Our results overall reveal a novel therapeutic approach to restore colonization resistance against *C. albicans* expansion by replacing Clostridia functionally with the drug 5-ASA. To describe this novel approach, we propose the term “faux-biotics” for chemical microbiota substitutes (e.g., 5-ASA) that replace probiotics (e.g., Clostridia) functionally. Whereas antibiotic prophylaxis interferes with engraftment of probiotics of fecal microbiota transplants, treatment with a faux-biotic offers the benefit that it is not inhibited or eliminated by antibiotics.

## MATERIALS AND METHODS

### Mice

For experiments using specific pathogen free (SPF) mice, 7-8 week-old female C57BL/6J mice were obtained from The Jackson Laboratory. Mice were maintained under SPF conditions throughout the duration of the experiment. For gnotobiotic experiments, age-, sex-, and strain-matched male and female C57BL/6J and Swiss-Webster mice, ages 6-10 weeks old, were bred in-house and were housed in Tecniplast Isocages throughout the duration of the experiments. Mice were fed autoclaved water and irradiated Teklad 2018 (2918) mouse chow.

To colonize the gnotobiotic mice with a cecal microbiota transfer (CMT) from either control or antibiotic-treated SPF mice, SPF mice were treated with 20 mg streptomycin vs. mock treatment and two days later cecal contents were collected, diluted at 1g/10 ml PBS, and stored in 25% glycerol at -80 C until use. Gnotobiotic mice were administered 200 μL of stored mock- or streptomycin-treated CMT by oral gavage. The CMT was given 1 week to colonize prior to further treatment. For the mice receiving first streptomycin-treated CMT followed by mock-treated CMT, mice were gavaged after 1 week with 200 μL of stored mock-treated CMT. For the C17 treatment group, mice were colonized with a mix of 17 human Clostridia isolates (Narushima, Sugiura et al. 2014) known to produce SCFAs. Strains were grown individually for 4 days in Eggerth-Gagnon broth, then equal volumes of broth from each strain were combined and 200 μL were administered by oral gavage to each mouse. For the 5-ASA treatment group, gnotobiotic mice were then placed on a diet containing 0.25% 5-ASA added to a base diet of Teklad 2018 mouse chow (Envigo). One week after the diet change or administration of C17, mice were infected with *C. albicans*.

At necropsy, mice were euthanized by overexposure to carbon dioxide followed by cervical dislocation. All procedures were approved by the UC Davis Animal Care and Use Committee.

### Infections

*Candida albicans* ATCC 28367 was streaked from a glycerol stock onto agar with 1% yeast extract, 2% peptone, 2% dextrose (YPD) + chloramphenicol at 100 ug/mL four days prior to infection, and re-streaked at two days prior to infection. To prepare the inoculum, colonies from the agar were resuspended in PBS, concentration was determined by OD600, and samples were diluted to the desired final concentration. Inocula were vortexed immediately prior to infection. Gnotobiotic mice were administered 10^5^ colony forming units (CFU) *C. albicans* in 100 μL phosphate buffered saline (PBS) by oral gavage. Specific pathogen free mice were given 10^5^ CFU (Fig. 3A, 5C, 6A-C) or 10^6^ CFU (Fig. 3B-C) *C. albicans* in 100 μL PBS by oral gavage, unless otherwise indicated. Inoculum CFUs were confirmed by plating on YPD agar + chloramphenicol to determine concentration.

One-2 days prior to or 1-5 days after infection with *C. albicans*, as indicated for each experiment, mice were randomized to receive 20 mg streptomycin in 100 μL sterile water by oral gavage or no treatment. For mice colonized with *C. albicans* prior to streptomycin treatment, colonization was confirmed, via measuring CFU/g feces, to be comparable between groups at the time of antibiotic administration.

*C. albicans* burden after infection was determined in feces and cecal contents by enumerating CFU on YPD agar + chloramphenicol. Prior to infection, mice were not colonized with any fungal organisms that grow on YPD agar. To collect feces, mice were placed in 70% ethanol-cleaned beakers cleaned until they produced a fecal pellet, which was then collected in 1 mL PBS, weighed, vortexed, and plated at 10-fold dilutions on YPD agar + chloramphenicol. Cecal contents collected at necropsy were similarly placed in PBS and plated. Plates were counted approximately 24 hours later. A presumptive identification of *C. albicans* was made for white, glistening, circular, smooth colonies growing on YPD agar containing chloramphenicol. Any colony growing on YPD agar + chloramphenicol with an atypical appearance was confirmed to be *Candida* spp. by Gram stain.

### SCFA analysis

Two fecal pellets per mouse were collected in 200 μL PBS. Samples were vortexed to disrupt particulate matter and then centrifuged at 6,000 rpm for 10 min to pellet any remaining debris. For each sample, 100 μL of supernatant was combined with 10 μL of a solution containing deuterated acetate, propionate, and butyrate so that each deuterated metabolite was at a final concentration of 100 μM. Samples were dried without heat in a vacuum dryer and then stored at -80°C until use.

Dried extracts were then solubilized by sonication in 0.1 mL anhydrous pyridine and then incubated for 20 min at 80°C. An equal amount of N-tert-butyldimethylsilyl-N-methyltrifluoroacetamide with 1% tert-butyldimethylchlorosilate (Sigma-Aldrich) was added, and the samples were incubated for 1 h at 80°C. Samples were centrifuged at 20,000 g for 1 min to remove leftover particles. One hundred microliters of the supernatant were transferred to an autosampler vial and analyzed by gas chromatography-mass spectrometry (Agilent 8890 Gas Chromatograph and Agilent 7000D Mass spectrometer). 1 μL of the sample was injected with a 1:50 split ratio at an injection temperature of 250°C on an HP 5ms Ultra Inert (2×15-m-length, 0.25-mm diameter, 0.25 μm film thickness) fused silica capillary column. Helium was used as the carrier gas with a constant flow of 1.2 mL/min. The gas chromatograph (GC) oven temperature started at 50°C for 5 min, rising to 110°C at 10°C/min and holding for 2 min, then raised to 310°C at 40°C/min with a final hold for 4 min. The interface was heated to 300°C. The ion source was used in electron ionization (EI) mode (70 V, 150 μA, 200°C). The dwell time for selected ion monitoring (SIM) events was 50ms. Both acetate, propionate and butyrate were quantified using SIM, with the monitored m/z, and experimentally determined retention times detailed in the following table:

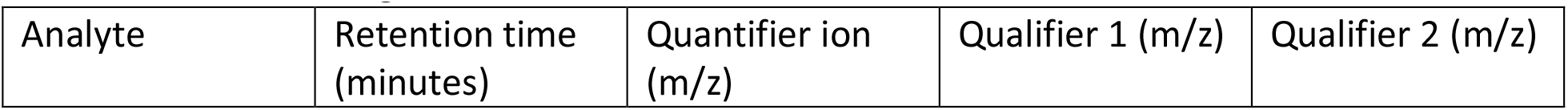
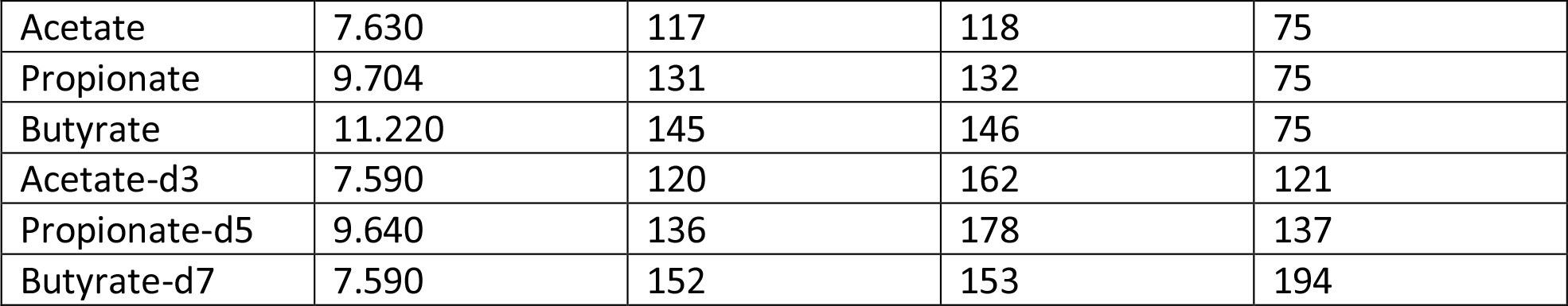

Efficient recovery of target metabolites was determined using deuterated compounds as internal standards. Quantification was based on external standards comprised of a series of dilutions of pure compounds, derivatized as described above at the same time as the samples.

### Histopathology

At necropsy, cecal tips were collected in individual histology cassettes and immediately placed in 10% phosphate-buffered formalin. After approximately 24 hours, cassettes were moved to 70% ethanol until processing. Samples were embedded in paraffin, mounted on slides, and stained with hematoxylin and eosin (H&E) by the UC Davis VMTH Pathology Lab.

Slides were scored in a blinded manner by a board-certified Veterinary Pathologist using standard scoring criteria (Supplemental Table 1) (Spiga, Winter et al. 2017).

### Colonocyte isolation and Real Time-PCR

Ceca and colons were opened and placed in PBS on ice after feces were gently removed.

Tissues were swirled in D-PBS until clean of fecal material, placed in DPBS with 0.03 M EDTA and 1.5 mM DTT on ice for 20 mins, then transferred to D-PBS with 0.03 M EDTA in 37°C for 10 min. Samples were shaken vigorously until colonocytes were released, remaining tissues were removed, and DPBS tubes containing colonocytes were spun at 800g for 5 min. at 4°C. After supernatant was removed, the colonocyte pellet was transferred to a cryovial and stored at - 80°C.

For mRNA isolation, colonocytes were thawed, TRI Reagent (Molecular Research Center) was added and incubated for 5 min., 200 μL chloroform was added, cells were spun at max speed, 4°C, for 15 min., and the aqueous phase was collected. An equal volume of 95% ethanol was added and samples were loaded onto Econo-Spin Columns (Epoch Life Science). Samples were washed with 3M sodium acetate, treated with PureLink DNase (Invitrogen), washed with 3x 70% ethanol+1% HEPES, and eluted with RNase-free water. RNA was quantified using the NanoDrop ND-1000 Spectrophotometer (Thermo Scientific). Complementary DNA was generated from 1 μg of RNA using MultiScribe reverse transcriptase (Applied Biosystems) with 10X RT PCR buffer (Applied Biosystems), 25mM MgCl2 (Applied Biosystems), dNTPs (Applied Biosystems), random hexamers (Applied Biosystems), and RNase inhibitor (Applied Biosystems). Samples were run on a PTC-200 Peltier Thermal Cycler (MJ Research) for 25°C (10 min.), 48°C (30 min.), 95°C (5 min.), then 4°C. Real-time PCR was performed with SYBR green (Applied Biosystems) and then primers listed in Supplemental Table 2 on a ViiA 7 real-time PCR system (Applied Biosystems) with the following cycling parameters: 50°C (2 min), 95°C (10 min), 40 cycles of 95°C (15 s) and 60°C (1 min). Results were analyzed using QuantiStudio Real-Time PCR software v1.3 (Applied Biosystems), and ΔΔ*C*_*T*_ was calculated with beta-actin as the control gene.

### Hypoxia staining and imaging

30-90 minutes prior to necropsy, mice were injected intraperitoneally with 100 mg/kg of pimonidazole HCl (Hypoxyprobe) in PBS. Staining with the Hypoxyprobe kit was done as previously described (Byndloss, Olsan et al. 2017, Cevallos, Lee et al. 2021). Briefly, paraffin-embedded tissues mounted on slides and prepared for staining with xylene (2x 10 min.) and ethanol (3 min. each in 95%, 80%, and 70%). Samples were treated with Proteinase K 20 mg/mL in TE buffer for 15 min. at 37°C, nonspecific binding was blocked with serum at room temperature for 1 hour, and slides were stained overnight at 4°C with the mouse IgG1 anti-PMDZ monoclonal antibody 4.3.11.3 (Hypoxyprobe). Slides were then stained with Cyanine3-labeled goat anti-mouse IgG (Jackson ImmunoResearch) for 90 min. at room temperature.

Between each staining step, slides were washed 3x 5 min. in PBS. Slides were briefly dried and mounted with Shandon Immu-Mount (Thermo Scientific).

Image numbers were randomized, blinded, and three representative images were collected on a Carl Zeiss AxioVision microcope with AxioVision 4.8.1 software (Zeiss) at 20X for scoring and 63X for image samples. Using ImageJ (NIH), the Texas Red channel (Cyanine3) was isolated, 3 representative slices of equal size containing the epithelial-lumen border were saved, and the Plot Profile for each was acquired. After unblinding, PMDZ peaks (epithelial-lumen borders) were aligned for each image and Profiles for the 9 slices associated with each mouse were averaged to obtain an average PMDZ Plot Profile and average PMDZ Peak for each mouse.

## Statistical analysis

Statistics were performed using GraphPad Prism 9.4.0 (GraphPad). An unpaired Student t test was performed on log-transformed CFU results, fold change in mRNA expression, and on short chain fatty acid levels. To compare peak pimonidazole staining intensities, a Mann-Whitney U test was performed. Throughout all figures, * = p<0.05, ** = p<0.005, *** = p<0.0005, and **** = p<0.00005.

## ACKNOWLEDGEMENTS

This work was supported by Microbiome Tri-institutional Partnership in Microbiome Research (TrIP) Seed funding award number 000582. Work in A.J.B.’s laboratory was supported by award 650976 from the Crohn’s and Colitis Foundation of America and by Public Health Service Grants AI044170, AI096528, AI112949, AI146432 and AI153069. This project was also supported by F32 AI161850 (H.P.S.). The project described was supported by the National Center for Advancing Translational Sciences, National Institutes of Health, through grant number UL1 TR000002 and linked award TL1 TR000133 (H.P.S.). The project described was supported by the National Center for Advancing Translational Sciences, National Institutes of Health, through grant number UL1 TR001860 (D.J.B.). The content is solely the responsibility of the authors and does not necessarily represent the official views of the NIH.

