## Supplemental Table 1 for "5-ASA can functionally replace Clostridia to prevent a post-antibiotic bloom of *Candida albicans* by maintaining epithelial hypoxia"

**Supplemental Table 1: Histopathology scoring**. Scoring criteria for H&E-stained cecal tips.

| **Score** | **Exudate** | **Epithelial damage** | **Neutrophils** | **Submucosal edema** |
| --- | --- | --- | --- | --- |
| 0 | Absent | Absent | 0-5 per hpf | Absent |
| 1 | Slight accumulation | Desquamation | 6-20 per hpf | Slight (<10%) |
| 2 | Mild accumulation | Mild erosion | 21-50 per hpf | Mild (10-20%) |
| 3 | Moderate accumulation | Marked erosion | 51-100 per hpf | Moderate (20-40%) |
| 4 | Severe accumulation | Ulceration | >100 per hpf | Severe (>40%) |
