## Supplemental Table 2 for "5-ASA can functionally replace Clostridia to prevent a post-antibiotic bloom of *Candida albicans* by maintaining epithelial hypoxia"

**Supplemental Table 2: Primers**.

| **Target genes** | **Name** | **Sequence** | **Reference** |
| --- | --- | --- | --- |
| *actb* | Beta actin | F: TGTCCACCTTCCAGCAGATGT  R: AGCTCAGTAACAGTCCGCCTAGA | Gu et al., 2013 |
| *lcn2* | Lipocalin 2 | F: ACATTTGTTCCAAGCTCCAGGGC  R: CATGGCGAACTGGTTGTAGTCCG | Godinez et al., 2008 |
| *tnf* | TNF-α | F: AGCCAGGAGGGAGAACAGAAAC  R: CCAGTGAGTGAAAGGGACAGAACC | Wilson et al., 2008 |
| *il17a* | IL-17A | F: AACCCCCACGTTTCTCAGCAAAC  R: GGACCCCTTTACACCTTCTTTTCATTG | Godinez et al., 2008 |
